# Sugar-phosphate metabolism regulates stationary phase entry and stalk elongation in *Caulobacter crescentus*

**DOI:** 10.1101/700559

**Authors:** Kevin D. de Young, Gabriele Stankeviciute, Eric A. Klein

## Abstract

Bacteria have a variety of mechanisms for adapting to environmental perturbations. Changes in oxygen availability result in a switch between aerobic and anaerobic respiration, whereas iron limitation may lead to siderophore secretion. In addition to metabolic adaptations, many organisms respond by altering their cell shape. *Caulobacter crescentus*, when grown under phosphate limiting conditions, dramatically elongates its polar stalk appendage. The stalk is hypothesized to facilitate phosphate uptake; however, the mechanistic details of stalk synthesis are not well characterized. We used a chemical mutagenesis approach to isolate and characterize stalk-deficient mutants, one of which had two mutations in the phosphomannose isomerase gene (*manA*) that were necessary and sufficient to inhibit stalk elongation. Transcription of the *pho* regulon was unaffected in the *manA* mutant; therefore, ManA plays a unique regulatory role in stalk synthesis. The mutant ManA had reduced enzymatic activity resulting in a 5-fold increase in the intracellular fructose 6-phosphate: mannose 6-phosphate ratio. This metabolic imbalance impaired the synthesis of cellular envelope components derived from mannose 6-phosphate, namely lipopolysaccharide O-antigen and exopolysaccharide. Furthermore, the *manA* mutations prevented *C. crescentus* cells from efficiently entering stationary phase. Deletion of the stationary-phase response regulator *spdR* inhibited stalk elongation in wild-type cells while overproduction of the alarmone ppGpp, which triggers growth arrest and stationary phase entry, increased stalk length in the *manA* mutant strain. These results demonstrate that sugar-phosphate metabolism regulates stalk elongation independently of phosphate starvation.

**Importance:** Bacteria have various mechanisms for adapting to environmental perturbations including morphological alterations. During phosphate limitation, *Caulobacter crescentus* dramatically elongates its polar stalk appendage. The stalk is hypothesized to facilitate phosphate uptake; however, the mechanism of stalk synthesis is not well characterized. We isolated stalk-deficient mutants, one of which had mutations in the phosphomannose isomerase gene (*manA*) that blocked stalk elongation, despite normal activation of the phosphate-starvation response. The mutant ManA produced an imbalance in sugar-phosphate concentrations that impaired the synthesis of cellular envelope components and prevented entry into stationary phase. Overproduction of the alarmone ppGpp, which promotes stationary phase entry, increased stalk length in the *manA* mutant demonstrating that sugar-phosphate metabolism regulates stalk elongation independently of phosphate starvation.

## Introduction

The diversity of bacterial cell shapes found in nature highlights the selective pressure for maintaining particular morphologies. The Gram-negative Alphaproteobacteria, *Caulobacter crescentus*, forms a unipolar stalk appendage during its asymmetric cell cycle. The dimorphic life cycle of *C. crescentus* produces one motile (swarmer) cell and one adherent (stalked) cell at each cell cycle (1). The swarmer cell has a polar flagellum and pili and is replication-incompetent. The swarmer then sheds its flagellum and, at the same pole, produces a holdfast - the strongest measured biological adhesive (2). After the holdfast is secreted, the stalk is formed and elongated from the holdfast pole, thereby causing the holdfast to be pushed away from the cell body and localized to the tip of the stalk. This stalked cell can perform DNA replication in preparation for cell division. During cell division, a new flagellum is synthesized at the opposite pole. As a result, following cytokinesis, the stalked cell maintains its stalk and immediately re-enters the cell cycle, while the other swarmer daughter cell is flagellated and enters a quiescent state from which it needs to emerge before synthesizing a new stalk and beginning a proliferative cycle.

In addition to its regulation by the cell cycle, stalk elongation is dramatically induced during phosphate limitation (3). Though the precise physiological functions of stalk elongation are not known, one proposed hypothesis was that the stalk acts as a “nutrient antenna” (4). Under the diffusive environment characteristic of freshwater lakes, nutrient flux is proportional to length; therefore, having a long thin appendage would be the most economical method of increasing cell length while minimizing surface area (5). A second proposed advantage of stalk elongation is that, in its natural environment, *C. crescentus* adheres to surfaces via holdfast at the stalk tip. By elongating the stalk, cells can rise above the surface to gain access to convective fluid flow thereby increasing nutrient availability (6).

While the timing of stalk elongation and its physiological consequences are fairly well understood, we know comparatively little about the mechanism of stalk synthesis. Phosphate limitation is known to activate the PhoRB two-component signaling pathway (3) and chromatin immunoprecipitation-DNA sequencing (ChIP-Seq) experiments identified nearly 50 genes regulated by PhoB in *C. crescentus* (7). Many of these genes are membrane transporters which is consistent with the stalk functioning as a nutrient scavenger.

The stalk is a true extension of the bacterial envelope containing inner- and outer-membranes as well as a peptidoglycan (PG) cell wall. The identification of PG synthesis enzymes responsible for stalk elongation has been elusive. PG synthesis during cell elongation and septation is performed by a family of mono- and bi-functional penicillin-binding proteins (PBPs) that have transglycosylase and/or transpeptidase activities. Deletion of the *C. crescentus* transglycosylases either individually or in combination does not prevent stalk formation in low phosphate conditions (8, 9); any of the paralogs (except PbpZ) suffice for cellular growth and stalk biogenesis. These data suggest that either the redundancy of this activity allows any PBP to synthesize stalk PG or there is a yet unidentified enzyme required for stalk PG insertion. Additionally, the stalk PG is enriched for LD-crosslinks (between *meso*-DAP residues on neighboring peptide stems) (10, 11); while these LD-crosslinks increase stalk PG resistance to lysozyme-mediated degradation (11), abrogation of LD-transpeptidation has no effect on stalk elongation (10, 11).

Studies using transposon mutagenesis or overexpression of fluorescent fusion proteins have successfully identified a number of stalk- or stalk-pole localized proteins, including: PbpC (12), BacA (12), StpX (13), BamE (14), and DipM (15). The deletion of these genes shortens, but does not eliminate, stalks in low-phosphate conditions. The difficulty in isolating a stalk-less *C. crescentus* strain implies that either: 1) stalk synthesis is an essential physiological process, 2) the synthesis enzymes have a secondary essential function and can therefore not be isolated by transposon mutagenesis, or 3) there is redundancy in the stalk synthesis pathway.

In this report, we used chemical mutagenesis to introduce single-nucleotide polymorphisms (SNPs) and screened for mutants with stalk elongation defects. We isolated a strain with mutations in *ccna_03732* (*manA*), a phosphomannose isomerase, that affected sugar-phosphate metabolism, cellular envelope biosynthesis, and entry into stationary phase. These physiological perturbations decreased stalk length despite normal induction of the *pho* regulon, suggesting that cellular metabolism regulates stalk elongation independently of phosphate starvation.

## Results

### Isolation of a stalk-deficient mutant

Genetic screens for phenotypes of interest are commonly performed by transposon mutagenesis. While this approach is quite powerful and allows for easy mapping of transposon insertions, it has the drawback that insertions in essential genes are highly unlikely since these mutants tend to be total loss-of-function and lethal. As an alternative approach, we used chemical mutagenesis to introduce SNPs into the *C. crescentus* genome and devised a screening methodology to isolate stalk-deficient mutants (see Materials and Methods). Briefly, *C. crescentus* cells grown in Hutner-Imidazole-Glucose-Glutamate media (HIGG) containing 1 mM phosphate (high phosphate) were treated with 1-methyl-3-nitro-1-nitrosoguanidine (NTG) to induce DNA mutations. The cells were washed and resuspended in HIGG-1 µM phosphate (low phosphate) to induce stalk elongation. To separate cells with stalk deficiencies, we reasoned that stalked cells float on top of a Percoll density gradient whereas non-stalked swarmer cells settle near the bottom of the gradient; therefore, we hypothesized that stalk-deficient cells would move to the bottom of a Percoll gradient. After 72 h of growth, cells were collected and subjected to three rounds of Percoll gradient centrifugation, each time collecting cells from the bottom of the gradient. Following the final gradient, cells were plated to isolate individual mutants. 384 individual colonies were grown in HIGG-1 µM phosphate and visually screened for stalk-phenotypes. One particular isolate, designated <u>S</u>talk-<u>D</u>eficient-<u>M</u>utant 1 (SDM1) had very short stalks compared to wild-type when grown in low phosphate (Fig. 1A).

**Figure 1.**
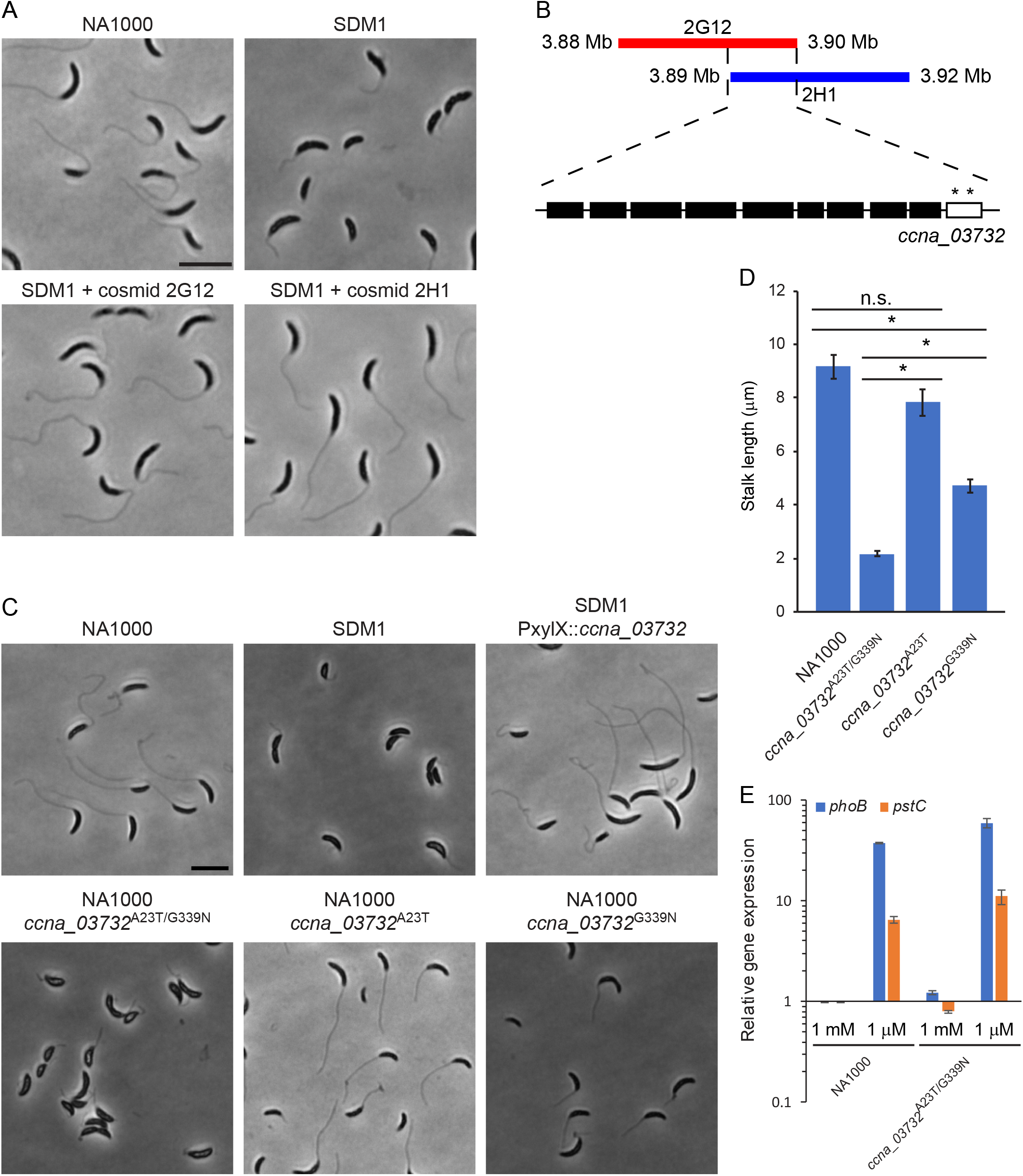
Isolation and mapping of a stalk-deficient mutant. (A) Screening a library of chemically-mutagenized *C. crescentus* strains identified <u>S</u>talk <u>D</u>eficient <u>M</u>utant 1 (SDM1). This mutant phenotype was rescued by complementation with two individual cosmids containing 20-30 kb genomic fragments. Scale bar: 5 µm. (B) A diagram of the two complementing cosmids shows their region of overlap. Within the overlapping genes, only *ccna_03732* has SNPs (denoted by *) as determined by whole-genome sequencing. (C-D) The SDM1 stalk phenotype was complemented by xylose-inducible expression of wild-type *ccna_03732*. Introduction of both *ccna_03732* SNPs into NA1000 was sufficient to produce the stalk-elongation defect, whereas each individual SNP had either no effect (A23T) or a partial effect (G339N). Scale bar: 5 µm. Stalk lengths were measured using ImageJ (error bars are SEM, ANOVA F(3,196)=76.8, P<0.0001; * post-hoc comparisons using Bonferroni test, P<0.05). (E) Wild-type and *ccna_03732*^A23T/G339N^ cells were grown for 24 h in HIGG-1 mM phosphate. For low phosphate samples, cells were washed twice in HIGG without phosphate, resuspended in HIGG without phosphate and grown for an additional 6 h. Total RNA was collected from each sample for qRT-PCR analysis of the PhoB-regulated genes *phoB* and *pstC*. Gene expression was normalized to *rpoD* and sample expressions were normalized to wild-type 1 mM phosphate (error bars are SEM, n=3). Strong induction of both genes confirmed that the *ccna_03732* SNPs do not affect the sensing of phosphate starvation.

### Mapping the SDM1 mutation

Whole-genome sequencing of the SDM1 strain yielded 79 potential SNPs. To identify the causative mutation for the stalk elongation phenotype, we performed complementation assays using a *C. crescentus* genomic cosmid library (16). Two cosmids, 2G12 and 2H1, were able to restore stalk elongation in low phosphate (Fig. 1A). These cosmids had an overlapping region of approximately 10 kb which contained 10 genes. Based on our sequencing data, of these 10 genes, only *ccna_03732* had any SNPs (Fig. 1B). The two missense mutations in *ccna_03732* resulted in amino acid substitutions A23T and G339N. To confirm that *ccna_03732* was necessary for the stalk elongation phenotype, we exogenously expressed wild-type *ccna_03732* in the SDM1 strain and observed recovery of stalk synthesis (Fig. 1C). Introduction of the two missense mutations into the *ccna_03732* chromosomal locus of wild-type *C. crescentus* phenocopied the SDM1 stalk deficiency, thereby demonstrating that these mutations were sufficient to inhibit stalk elongation (Fig. 1C). To determine which SNP was responsible for the SDM1 phenotype, we introduced each SNP individually into the *C. crescentus* chromosome; the A23T mutation had no significant effect on stalk elongation whereas the G339N mutation only partially recapitulated the elongation defect seen in SDM1 (Fig. 1C-D). Therefore, we conclude that the SDM1 stalk-elongation phenotype required both SNPs. The *ccna_03732* gene is predicted to be essential (17) and we were unable to generate a deletion of this gene using standard allelic replacement methods, thereby demonstrating the power of chemical mutagenesis for phenotypic screens to isolate mutations leading to altered function rather than total loss-of-function. We confirmed that the *ccna_03732* mutations did not affect the ability of *C. crescentus* to sense low-phosphate concentrations; phosphate starvation similarly induced the expression of PhoB-regulon genes *phoB* and *pstC* in wild-type and mutant cells (Fig. 1E). Thus, *ccna_03732* affects stalk synthesis independently of phosphate-mediated regulation.

### Determining the enzymatic function of CCNA_03732

CCNA_03732 is annotated as a YihS-domain containing epimerase (GenBank ACL97197.3) and a BLAST search identified YihS as the closest *Escherichia coli* homologue (18). Exogenous expression of *E. coli* YihS in the *ccna_03732*^A23T/G339N^ background did not complement the stalk defect (Fig. 2A) despite being well expressed (Fig. 2B). YihS is part of a larger family of N-acyl-D-glucosamine 2-epimerases (AGEs) which isomerize a wide variety of carbohydrate substrates (19). *E. coli* encodes three additional AGE proteins (RffE, NanE, and ManA); therefore, we tested whether any of these genes could complement the CCNA_03732 mutation. Only expression of ManA was able to rescue stalk elongation (Fig. 2A); therefore, we will refer to CCNA_03732 as ManA. ManA is a phosphomannose isomerase (PMI) which catalyzes the interconversion of mannose 6-phosphate (M6P) and fructose 6-phosphate (F6P) (20). Although *E. coli* ManA expression does not fully recover stalk elongation when compared to wild-type (Fig. 2A), this may because *E. coli* ManA is a type-I PMI whereas *C. crescentus* ManA appears to be a type-III enzyme (21). The three subfamilies of PMIs have similar enzymatic functions but little sequence homology (22).

**Figure 2.**
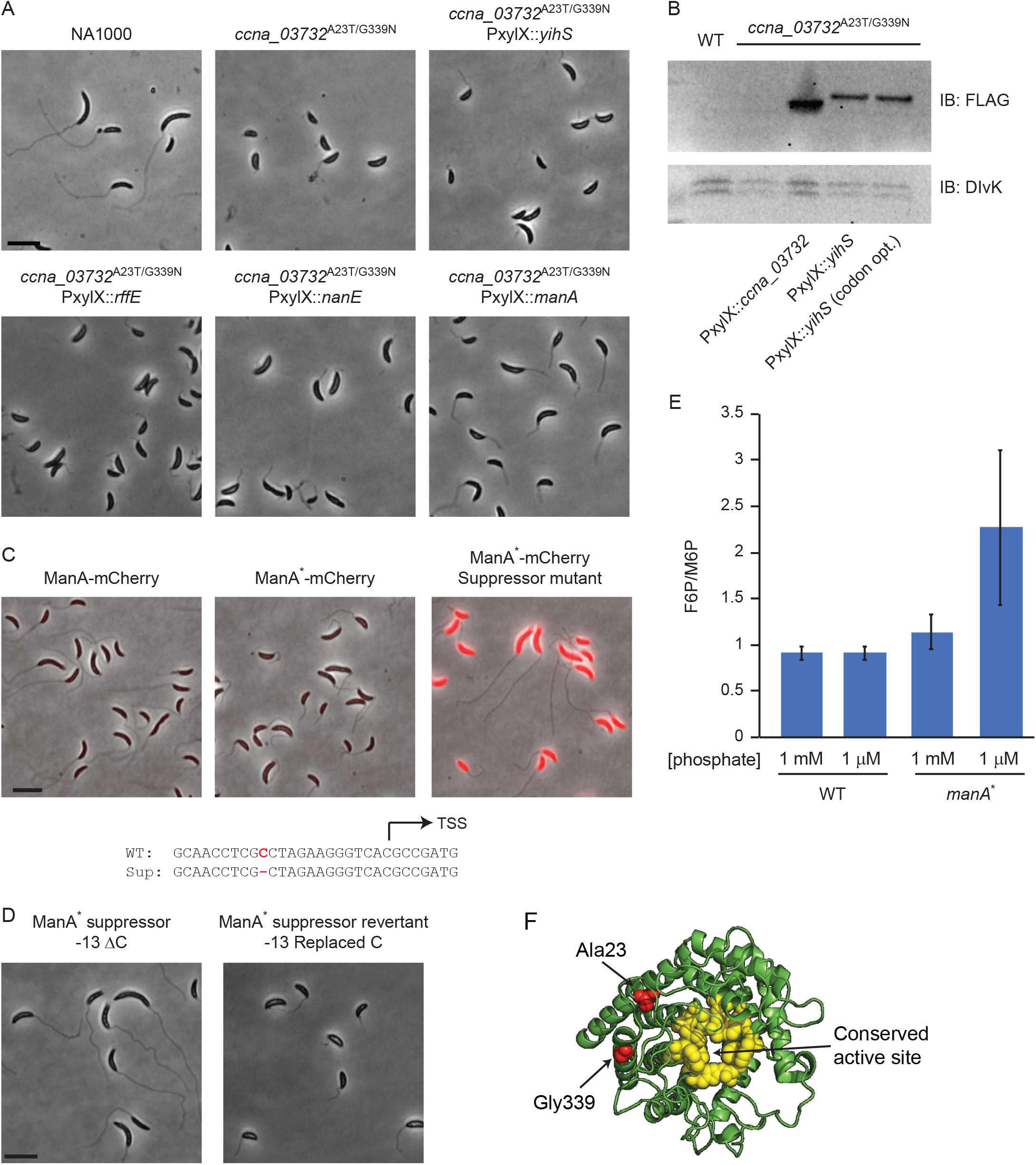
CCNA_03732 is a phosphomannose isomerase. (A) Exogenous expression of the closest *E. coli* homologue to CCNA_03732, YihS, did not complement the stalk elongation defect. Screening the other AGE-family isomerases demonstrated that only ManA can rescue stalk synthesis in the *ccna_03732*^A23T/G339N^ strain. (B) Western blot analysis of FLAG-tagged CCNA_03732 and YihS (both wild-type and codon-optimized) showed that they are expressed at comparable levels. Therefore, the inability of YihS to rescue stalk-elongation is a reflection of differing enzymatic activities rather than expression level. (C) Native-locus mCherry fusions to ManA showed that the stalk elongation phenotype can be suppressed by dramatic overexpression of ManA^*^. Sequencing of the *manA* locus in the suppressor strain identified a single base deletion upstream of the *manA* transcription start site. Scale bar: 5 µm. (D) Reinsertion of the deleted base into the suppressor strain reverted the stalk deficiency phenotype. Scale bar: 5 µm. (E) Metabolomic analysis of F6P and M6P levels showed that wild-type cells maintain a 1:1 ratio in both low and high phosphate, whereas *manA*^*^ cells increase their relative levels of F6P as phosphate decreases (error bars are SEM, n=3). (F) A homology model of ManA (based on PDB 1FP3) shows that the ManA mutations (red) are near one another but not near the enzyme active site (yellow).

To investigate the mechanism of the *ccna_03732*^A23T/G339N^ (referred to as *manA*^*^) mutations, we performed a suppressor screen to isolate mutants which regained stalk elongation (see Materials and Methods). This screen yielded a suppressor with a one-base deletion at the - 13-position relative to the transcription start site (Fig. 2C) leading to overexpression of ManA (Fig. 2C). Replacing the deleted cytosine in the suppressor mutant caused a reversion to the short-stalk phenotype demonstrating the sufficiency of this SNP to rescue stalk elongation (Fig. 2D). Since overexpression of ManA restored stalk synthesis, we concluded that the ManA^*^ mutant is a hypomorph with reduced enzymatic activity. Comparison of F6P/M6P ratios in wild-type and *manA*^*^ strains demonstrated that wild-type cells maintained a 1:1 ratio in both high and low phosphate whereas the ratio rises to 2:1 for *manA*^*^ cells in low phosphate (Fig. 2E). These data suggest that in low phosphate, ManA^*^ cannot efficiently convert F6P to M6P. Mapping the A23T and G339N mutations onto a homology model of ManA (produced using Phyre 2.0 (23) based on PDB structure 1FP3 (24)) showed that the mutations are positioned near one another yet distant from the predicted protein active site (Fig. 2F) (19) making it difficult to propose a mechanism for reduced ManA activity.

### Effects of ManA^*^ on bacterial envelope synthesis

Metabolic network maps (KEGG: amino sugar and nucleotide sugar metabolism) place ManA in a central position for regulating bacterial envelope synthesis (Fig. 3A) (25). F6P is a precursor for N-acetylglucosamine and N-acetylmuramic acid which are important metabolites for PG and core lipopolysaccharide (LPS) synthesis. M6P, which is metabolized to GDP-mannose, serves as a precursor for LPS O-antigen and exopolysaccharide (EPS) synthesis (26). Analysis of LPS demonstrated that the *manA*^*^ cells have approximately 83% less O-antigen-containing smooth-LPS relative to wild-type cells, while rough LPS (lacking O-antigen) was unaffected (Fig. 3B); disruption of *wbqP* inhibits O-antigen synthesis, likely by preventing the attachment of the first sugar (perosamine) to the undecaprenol carrier lipid, and is included as a negative control (27, 28). Comparison of the muropeptide content of PG from wild-type and *manA*^*^ cells did not reveal any gross changes in PG composition (Fig. 3C). Lastly, production of EPS was assessed by the presence of a mucoid phenotype when grown on agar plates supplemented with 3% sucrose; wild-type cells producing EPS were mucoid while *manA*^*^ cells were dull in appearance (Fig. 3D). Non-EPS producing strains (CB15 and NA1000 ΔMGE) were included as controls (26). Together, these findings showed that F6P-dependent metabolism was unaffected in the *manA*^*^ strain, whereas M6P-dependent processes were inhibited. This is consistent with the ManA^*^ mutant being an enzymatic hypomorph leading to an increased F6P:M6P ratio (Fig. 2C, E). To determine whether these changes in cellular envelope were the cause of the stalk elongation defect, stalk lengths were measured in NA1000 strains lacking either EPS (ΔMGE) or O-antigen (Tn5::*wbqP* (27)) (Fig. 3E). Inhibition of EPS production had no significant effect on stalk length; however, O-antigen disruption reduced stalk length although not to the same degree as the *manA*^*^ strain (Fig. 3E). While the partial effect of Tn5::*wbqP* on stalk length suggests that O-antigen synthesis plays a role in stalk elongation, an alternative explanation is that the inhibition of this pathway alters the flux through connected metabolic pathways leading to stalk elongation defects.

**Figure 3.**
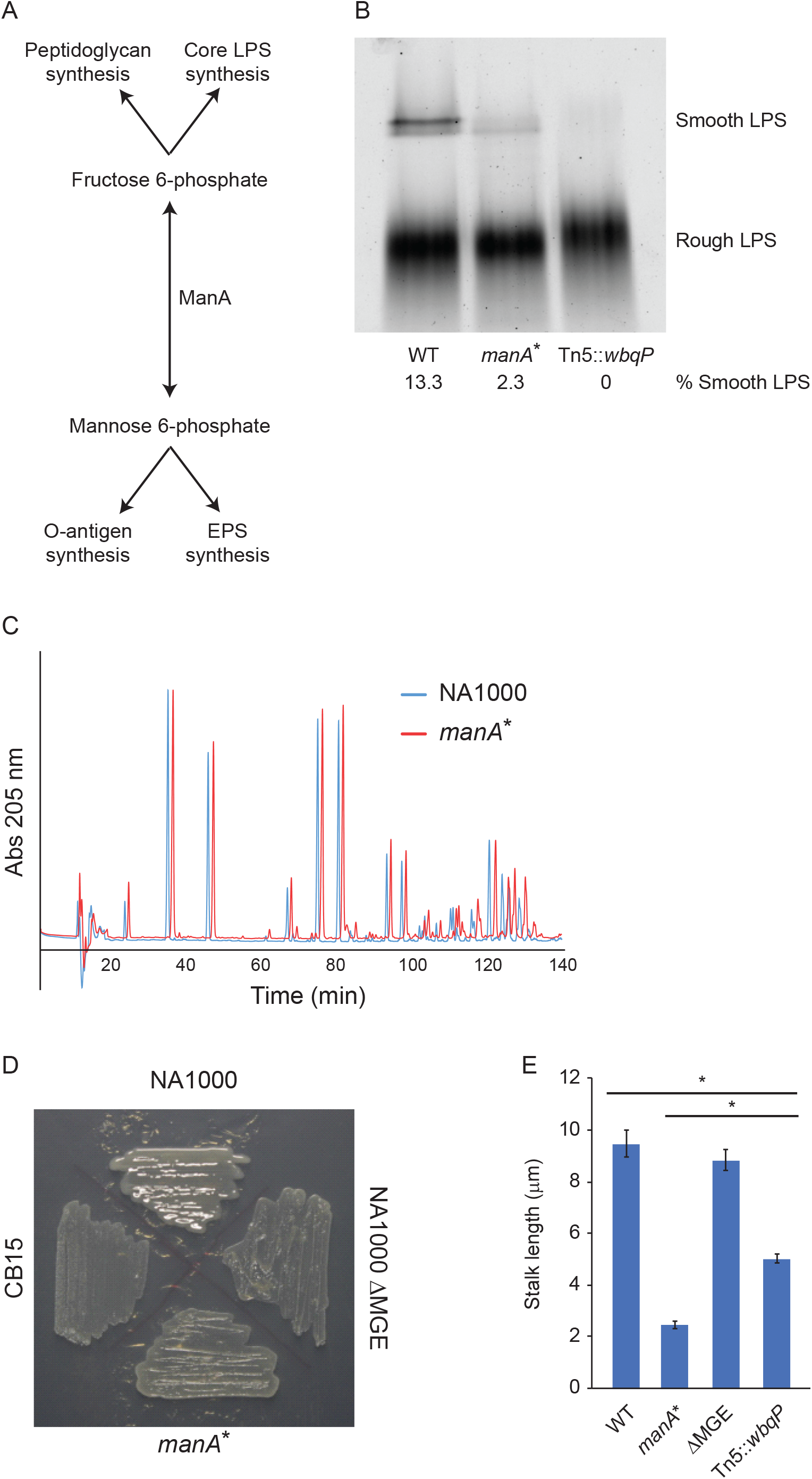
ManA^*^ results in alterations to specific cellular envelope components. (A) ManA regulates the interconversion of F6P and M6P. These metabolites are important precursors for the synthesis of cellular envelope components including LPS, PG, and EPS. (B) LPS was extracted from wild-type, *manA*^*^, and Tn5::*wbqP* cells grown in the HIGG-1 µM phosphate and separated by SDS-PAGE. The *manA*^*^ strain produced wild-type levels of rough-LPS but diminished smooth-LPS. The Tn5::*wbqP* cells do not produce any O-antigen. The percentage of smooth LPS was determined from the band intensities of the smooth and rough LPS for each sample. (C) Comparisons of muropeptides from wild-type and *manA*^*^ cells grown in HIGG-1 µM phosphate did not yield any gross changes in PG composition. (D) The indicated strains were streaked onto HIGG plates supplemented with 3% sucrose to induce EPS production and mucoidy. Wild-type cells had a distinct mucoid appearance whereas *manA*^*^ was matte in appearance signifying a decrease in EPS production. CB15 and NA1000 ΔMGE were non-EPS-producing control strains. (E) The indicated strains were grown in HIGG-1 µM phosphate for 48 h and stalk lengths were measured using ImageJ (error bars are SEM, ANOVA F(3,218)=74.6, P<0.0001; * post-hoc comparisons using Bonferroni test, P<0.05). The Tn5::*wbqP* strain, which does not make O-antigen, had a partial effect on stalk elongation compared to the *manA*^*^ strain.

### ManA regulates entry into stationary phase

Since F6P and M6P feed into a variety of metabolic pathways, we performed metabolomic analyses of wild-type and *manA*^*^ cells grown in low-phosphate to assess global changes in cellular metabolism. As expected, wild-type cells had higher levels of M6P as compared to *manA*^*^ cells; additionally, we found higher levels of TCA cycle intermediates in wild-type cells (Fig. 4A). Accumulation of TCA cycle intermediates is consistent with bacteria entering stationary phase (29, 30), therefore, we performed long term growth curves to assess stationary phase entry. When cultured in HIGG-30 µM phosphate, wild-type growth plateaued at OD_660_=1.8; by contrast, *manA*^*^ cells continued to divide to OD_660_=2.5, suggesting that they failed to enter stationary phase (Fig. 4B). As observed with regards to stalk elongation, exogenous expression of *manA* in the mutant background restored wild-type growth kinetics (Fig. 4B). Consistent with their increased proliferation, *manA*^*^ cells had lower expression of known stationary-phase genes *cspD*, *katG*, and *spdR* (31) following 48 h of culture in low-phosphate media (Fig. 4C) and deletion of the stationary-phase response regulator *spdR* inhibited stalk elongation (Fig. 4D).

**Figure 4.**
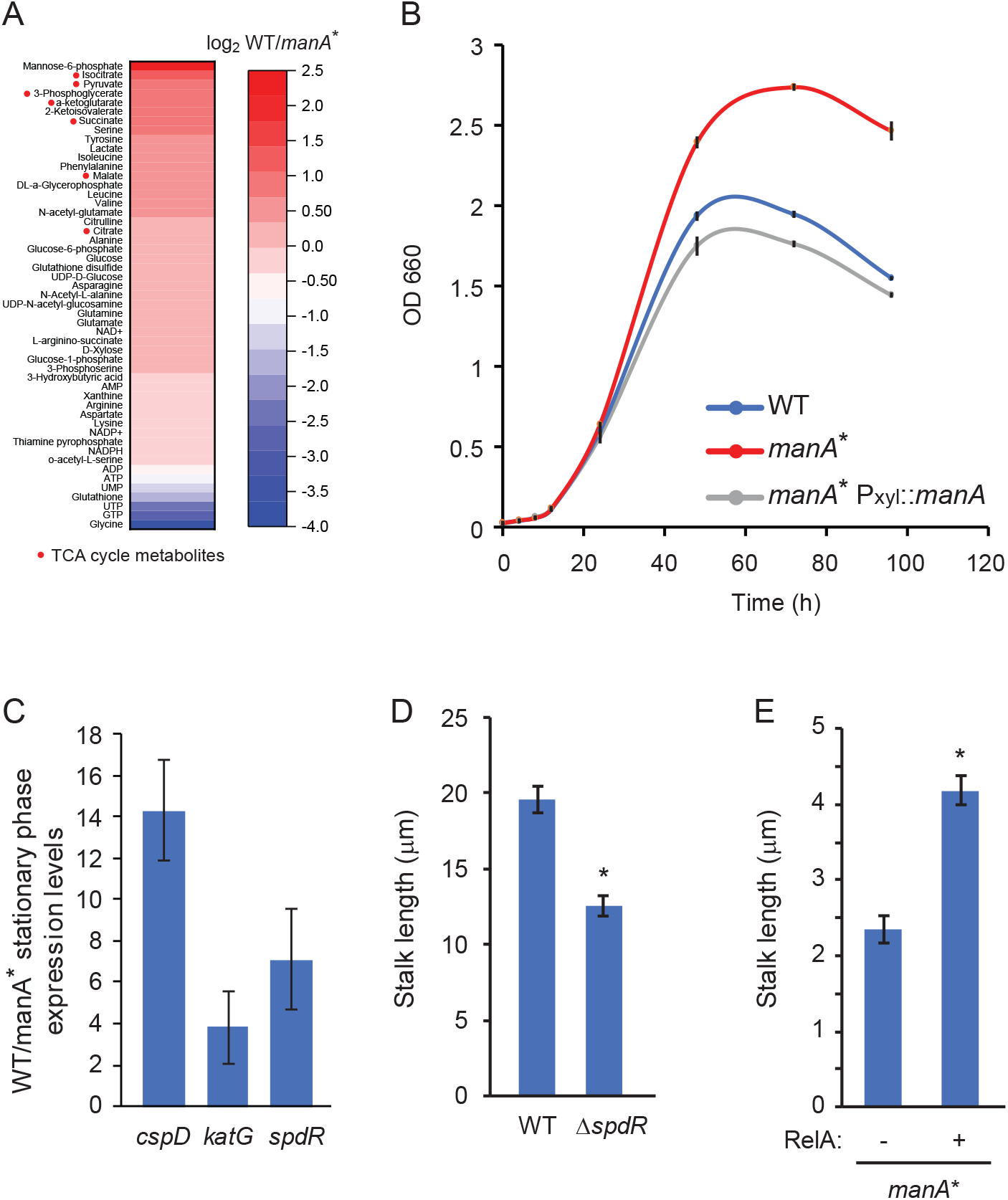
*manA*^*^ cells have a defect in stationary-phase entry which contributes to the stalk-elongation phenotype. (A) Metabolomic analyses of wild-type and *manA*^*^ cells grown in HIGG-1 µM phosphate demonstrated that wild-type cells have higher levels of M6P as well as TCA cycle metabolites (red circles) (n=3). (B) Growth curves of wild-type, *manA*^*^, and complemented *manA*^*^ cells in HIGG-30 µM phosphate showed that *manA*^*^ grows to a higher OD than wild-type (error bars are SEM, n=3). (C) Wild-type and *manA*^*^ cells were grown in HIGG-30 μM phosphate for 48 h. Wild-type and *manA\** cultures were back diluted into HIGG-30 μM phosphate to an OD_660_ = 0.03 and allowed to grow for 48 h. Expression of the indicated stationary-phase genes was measured by qRT-PCR and relative mRNA levels between wild-type and *manA*^*^ were quantified. Gene expression was normalized to *rpoH* mRNA (error bars are SEM, n=3). (D) Wild-type and Δ*spdR* cells were grown for 48 h in HIGG-1 µM phosphate and stalk lengths were measured. *spdR* deletion inhibits stalk elongation (error bars are SEM, * Mann-Whitney *U*=2778, *n_1_*=58, *n_2_*=61, *P* < 0.05 two-tailed). (E) *manA*^*^ cells with or without exogenous expression of constitutively-active RelA were grown in HIGG-1 µM phosphate with 0.003% xylose and stalk length was measured by microscopy. RelA expression rescued stalk elongation (error bars are SEM, * Mann-Whitney *U*=1313, *n_1_*=70, *n_2_*=87, *P* < 0.05 two-tailed).

Stationary-phase entry and the stringent response are, in part, regulated by the production of the alarmone guanosine 3’,5’-bispyrophosphate (ppGpp) (32, 33). Overexpression of the N-terminal domain of the ppGpp synthetase RelA results in the constitutive production of ppGpp and decreased growth rate in *E. coli* (34). To test whether stationary phase entry was necessary for stalk elongation in *C. crescentus*, we overexpressed truncated RelA in the *manA*^*^ cells; overproduction of ppGpp resulted in a significant increase in stalk length (Fig. 4E).

## Discussion

*C. crescentus* stalks elongate under phosphate-limited conditions, however the mechanism is not well understood. Perturbation of genes associated with peptidoglycan synthesis results in shorter stalks (8, 9, 12); however, this system appears to be highly redundant as even the deletion of multiple PBPs does not abrogate stalk synthesis completely. Recently, a fluorescent fusion to the cytoskeletal protein MreB was shown to be functional with regards to cell growth, but this fusion did not localize at the poles nor support stalk formation (10), suggesting that the GFP-tag interferes with protein-protein interactions necessary for stalk synthesis.

To identify novel regulatory mechanisms of stalk elongation, we used chemical mutagenesis to introduce single-nucleotide polymorphisms (SNPs) and isolated mutants with stalk elongation defects. One particular strain had two SNPs in *ccna_03732* (*manA*), a phosphomannose isomerase (Fig. 2A), both of which were required for a stalk-deficient phenotype (Fig. 1A-D). These SNPs affected the relative levels of F6P and M6P (Fig. 2E) and cellular envelope biosynthesis (Fig. 3A-D), but not the transcriptional response to phosphate limitation (Fig. 1E). These findings demonstrate that cellular metabolism regulates stalk elongation independently of phosphate starvation.

Disruption of sugar-phosphate metabolism results in cell shape phenotypes in a wide range of organisms. In *Bacillus subtilis*, deletion of *manA* leads to cells with an elongated spheroid morphology (35). This morphology is due, in part, to a decrease in wall techoic acid (WTA) production which affects cell wall architecture (35). PMI plays a role in eukaryotic cell shape as well. Deletion of PMI in the fungus *Aspergillus fumigatus* affects conidiation and produces a thickened chitin-rich cell wall (36), while in the protozoan *Leishmania mexicana*, Δ*pmi* promastigotes were shorter and rounder than wild-type (37).

While *manA* disruption affects cell shape in both *C. crescentus* and *B. subtilis*, it appears that the mechanisms may be quite different. In Gram-positive *B. subtilis*, ManA specifically reduces the levels of the cell wall carbohydrates glucose and N-acetylgalactosamine (GalNAc) without affecting N-acetylglucosamine (GlcNAc). Thus, Δ*manA* affects WTA but not PG synthesis. By contrast, Gram-negative *C. crescentus* does not produce WTA, and the composition of the PG was not affected in the *manA*^*^ strain (Fig. 3C); thus, another mechanism is required to explain the stalk elongation phenotype. While the *manA*^*^ mutations affected LPS O-antigen and EPS production, these envelope perturbations were not solely responsible for the mutant cell defect (Fig. 3E).

In addition to interfering with cellular envelope biosynthesis, the metabolic imbalance produced by the *manA*^*^ mutations prevents stationary-phase entry of *C. crescentus* (Fig. 4A-C). The accumulation of sugar-phosphates triggers a variety of bacterial stress responses. In *B. subtilis*, a build-up of M6P leads to increased expression of the stress sigma factors σ^X^ and σ^W^ (38) as well as derepression of the *glcR-phoC* operon (39). In *E. coli*, elevated concentrations of sugar-phosphates leads to growth inhibition (40), induction of the stringent response (41), and expression of the stress-response gene *uspA* (42). This response is mediated, in part, by the small RNA *sgrS* which directs the degradation of mRNAs important for sugar-phosphate transport including *ptsG* and *manXYZ* as well as the stabilization of the phosphatase *yigL* (43–45). Interestingly, while alterations to sugar-phosphate homeostasis promote the stringent response and growth arrest in *B. subtilis* and *E. coli*, F6P accumulation in *C. crescentus* due to the *manA*^*^ mutations prevents stationary-phase entry. *C. crescentus* does not encode homologues of SgrRS, UspA, or GlcR; therefore, further investigations will be required to identify the sensor and downstream targets of sugar-phosphate stress in *C. crescentus*.

## Materials and Methods

### Bacterial strains, plasmids, and growth conditions

The strains, plasmids, and primers used in this study are described in Supplemental Tables S1, S2, and S3, respectively. Details regarding strain construction are available in the Supplementary Methods.

*C. crescentus* wild-type strains NA1000, CB15, and their derivatives were grown at 30 °C in peptone-yeast-extract (PYE) medium (46) for routine culturing. To regulate phosphate levels, *C. crescentus* was grown in Hutner-Imidazole-Glucose-Glutamate (HIGG) media with varying concentrations of phosphate (1-1000 µM) (47). *E. coli* strains were grown at 37 °C in LB medium. When necessary, antibiotics were added at the following concentrations: kanamycin 30 µg ml^−1^ in broth and 50 µg ml^−1^ in agar (abbreviated 30:50) for *E. coli* and 5:25 for *C. crescentus*; tetracycline 12:12 *E. coli* and 1:2 *C. crescentus*. Gene expression was induced in *C. crescentus* with 0.003-0.3% (w/v) xylose. Growth curves were determined by measuring OD (660 nm) at the indicated time points.

### Chemical mutagenesis and screening for stalk-elongation mutants

*C. crescentus* cells (1 mL) were grown in HIGG-1 mM phosphate overnight and treated with 200 µg mL^−1^ 1-methyl-3-nitro-1-nitrosoguanidine (NTG, TCI America) for 50 minutes at 30 °C to induce DNA mutations. The cells were washed once in HIGG without phosphate and resuspended in 100 mL HIGG-1 µM phosphate to promote stalk elongation. After 72 h, cells were pelleted at 16,000 x g for 20 min, resuspended in 5.5 mL of HIGG-1 µM phosphate, and mixed 1:1 with Percoll (GE Healthcare) to establish a density gradient. This mixture was centrifuged for 1 h at 9,500 x g and cells from the bottom of the tube were recovered using a Pasteur pipette. After two additional Percoll gradients, the bottom cells were plated on PYE plates to isolate individual mutants. 384 individual colonies were grown in HIGG-1 µM phosphate and visually screened for stalk defects.

### Whole-genome sequencing

Genomic DNA was isolated from NA1000 and SDM1, barcoded sequencing libraries were constructed using the NEBNext Fast DNA Fragmentation & Library Prep Set for Ion Torrent (New England Biolabs), and templated using the Ion PI Hi-Q Chef kit (Thermo Fisher). The libraries were sequenced on an Ion Proton system (Thermo Fisher) (NA1000: 2,148,749 reads, mean read length=132 bp, mean sequencing depth=70X; SDM1: 1,976,601 reads, mean read length=132 bp, mean sequencing depth=64X). Reads were mapped to the NA1000 genomic sequence (NCBI NC_011916.1) and variants were called using Ion Reporter software (Thermo Fisher). SNPs present in both SDM1 and our lab strain of NA1000 were eliminated from analysis.

### Cosmid conjugation

*E. coli* cosmid-library strains (kindly provided by Lucy Shapiro, Stanford University) (16) and SDM1 *C. crescentus* cells were grown overnight in LB and PYE, respectively. 100 µL of each cosmid-strain was mixed with 900 µL SDM1, centrifuged at 8,000 x *g* for 2 min, washed once in PYE, and the final pellet was resuspended in 20 µL PYE. The mixed and concentrated samples were dropped onto PYE plates and incubated at 30 °C for 6 h. Samples of the bacterial conjugants were streaked onto PYE plates containing 25 µg mL^−1^ kanamycin and 30 µg mL^−1^ nalidixic acid; *C. crescentus* cells are naturally resistant to nalidixic acid allowing for selection of *C. crescentus* cells that have received the cosmid.

### *manA*^*^ suppressor screen

A single colony of the *manA*^*^ strain grown 2mL of HIGG-1mM phosphate overnight at 30□. 1 mL of the culture was washed twice in HIGG (without phosphate), resuspended in HIGG-1μM phosphate, and grown for 2 days at 30□. The culture was centrifuged at 10,000 x g for 10 minutes. The supernatant containing buoyant potential suppressor cells was collected and centrifuged at 28,000 x g for 30 minutes. The pelleted cells were then resuspended in 20mL of HIGG-1μM phosphate. This purification process was repeated every 48 h; after two weeks the culture was streaked onto PYE plates to isolate individual colonies. Colonies were subsequently grown in HIGG-1μM phosphate and screened by microscopy for stalk recovery.

### Microscopy and image analysis

Cells were spotted onto 1% agarose pads made in the corresponding growth medium. Microscopy was performed on a Nikon TiE inverted microscope equipped with a Prior Lumen 220PRO illumination system, Zyla sCMOS 5.5-megapixel camera, CFI Plan Apochromat 100X oil immersion objective (NA 1.45, WD 0.13 mm), and NIS Elements software for image acquisition. Stalk lengths were measured using ImageJ v. 1.48q (NIH).

### Quantitative RT-PCR (qRT-PCR)

RNA was extracted from bacterial cultures using the Qiagen RNeasy kit. Following DNase digestion, RNA (5 ng µl^−1^) was reverse transcribed using the High Capacity cDNA Reverse Transcription Kit (Applied Biosystems). 1 µl of cDNA was used as a template in a 10 µl qRT-PCR reaction performed with Power SYBR reagent (Applied Biosystems). qRT-PCR was performed on an ABI QuantStudio 6 using the ΔΔCt method. *rpoD* or *rpoH* expression was used as the loading control as indicated.

### Western blot analysis

*C. crescentus* cells were grown overnight in the indicated medium and cell concentration was normalized by OD_660_. 1 mL samples of OD_660_=0.5 were collected by centrifugation (8,000 x *g*, 2 min), resuspended in 100 µL 1X Laemmli buffer, and boiled for 5 min to denature proteins. Protein samples were separated on 10% SDS-PAGE gels, transferred to PVDF membrane, and probed using the following antibodies: FLAG (1:500, sc-166355, Santa Cruz) and DivK (1:1000, kind gift from Lucy Shapiro, Stanford University (48)).

### Metabolomics

*C. crescentus* cells were grown overnight in HIGG-1 mM phosphate, back diluted, and grown to OD_660_=0.3. For high-phosphate samples, 5 mL of OD_660_=0.3 was filtered onto a 0.2 µm pore-size nylon membrane (Millipore GNWP04700). For low-phosphate samples, 5 mL of OD_660_=0.3 was washed twice in HIGG lacking phosphate, resuspended in 5 mL HIGG-1 µM phosphate, grown for 6 h, and then filtered onto nylon membranes. The filters were immediately quenched in 1.2 mL ice-cold 40:40:20 acetonitrile:methanol:water containing 0.5% (v/v) formic acid. The samples were incubated at −20 °C for 15 minutes and the solutions were transferred to pre-chilled 2 mL microcentrifuge tubes containing 50 mg 0.1 mm glass beads (Med Supply Partners, NA-GB01-RNA). The solvent was neutralized by adding 100 µL 1.9M ammonium carbonate and cells were lysed on a Qiagen TissueLyser for 5 min at 30 Hz. Samples were centrifuged at 16,000 x *g* for 10 min at 4 °C to pellet unbroken cells and the supernatant was transferred to a pre-chilled microcentrifuge tube and frozen on dry ice. Samples were sent to the Metabolomics Core Facility of the Cancer Institute of New Jersey (New Brunswick, NJ) for analysis by liquid chromatography/mass-spectrometry (LC/MS). Since F6P and glucose 1-phosphate have the same molecular weight and co-elute by LC, it is impossible to distinguish between these two sugar-phosphates. To enable quantification of F6P, we replaced the glucose in our standard HIGG media with deuterated 2-D glucose (Cambridge Isotope Laboratories, DLM-1271). During the isomerization of glucose to fructose, the deuterium at C-2 is replaced with hydrogen, leading to a 1 mass-unit shift between these sugars and enabling their distinction by MS.

### Lipopolysaccharide (LPS) purification and analysis

LPS was purified essentially as previously described (28, 49). Briefly, 5 ml of *C. crescentus* cells grown in HIGG-1 µM phosphate (OD_660_ = 0.5) were collected and washed once in 10 mM HEPES, pH 7.2. Cells were resuspended in 250 µl TE buffer (10 mM Tris, 1 mM EDTA, pH 7.2) and frozen overnight at −20 °C. Cells were thawed, treated with 1 µl DNase (0.5 mg ml^−1^), 20 µl lysozyme (10 mg ml^−1^), and 3 µl MgCl_2_ (1M), and incubated at room temperature for 15 minutes. For each sample, 36.25 µl was mixed with 12.5 µl 4X SDS-sample buffer and boiled at 100 °C for 10 minutes. After cooling to room temperature, 1.25 µl proteinase K (20 mg ml^−1^) was added and samples were incubated at 60 °C for 1 hour. LPS samples were resolved on a 12% SDS-PAGE gel and stained with the Pro-Q Emerald 300 LPS stain kit according to the manufacturer’s protocol (Thermo Scientific). Images were acquired on a Bio-Rad ChemiDoc MP using UV excitation and a 530 nm emission filter.

### Peptidoglycan (PG) purification and analysis

*C. crescentus* cells (500 mL) were grown in HIGG-1 µM phosphate. Peptidoglycan muropeptides were purified from *C. crescentus* as previously described (50) and separated on a reversed-phase C18 column (Thermo Scientific; 250 x 4.6-mm column, 3-µm particle size) held at 55 °C. The LC solvent system consisted of 50 mM sodium phosphate [pH 4.35] with 0.4% sodium azide (solvent A) and 75 mM sodium phosphate, pH 4.95 + 15% (v/v) methanol (solvent B). The solvent flow rate was 0.5 mL min^−1^ and a linear gradient to 100% solvent B was performed over 135 min. Muropeptide elution was monitored at 205 nm.

### Exopolysaccharide (EPS) production assay

Strains were streaked onto HIGG-1 µM phosphate agar plates supplemented with 3% (w/v) sucrose. EPS production was determined by assessing mucoidy. CB15 and NA1000 ΔMGE were used as non-EPS producing controls (26).

## Acknowledgements

We thank Lucy Shapiro (Stanford University) and Christine Jacobs-Wagner (Yale University) for providing reagents and Xiaoyang Su and the metabolomics core facility at the Cancer Institute of New Jersey for their experimental assistance. Funding was provided by National Science Foundation CAREER Award MCB-1553004 to E.A.K.

