## Supplementary material for "Sugar-phosphate metabolism regulates stationary phase entry and stalk elongation in *Caulobacter crescentus*": Table S

Supplementary methods

**Strain construction.** Creation of the various *C. crescentus* chromosomal point-mutation and deletion strains was done by double-homologous recombination using both positive and negative selection. The suicide plasmid pNPTS138 has both a kanamycin resistance cassette as well as the *sacB* gene which is toxic in the presence of sucrose. Genomic fragments (1000 bp) either centered on or flanking the chromosomal region of interest were ligated into the pNPTS138 vector. The plasmid was transformed into *C. crescentus* and recombinants were selected on kanamycin plates. Individual colonies were grown overnight in PYE (without antibiotic) and streaked out onto PYE-3% sucrose plates to recover colonies that performed the second recombination. Colonies were screened for the gene mutation or deletion by PCR and streaked out onto plain PYE and PYE-kanamycin plates to confirm the loss of the plasmid backbone.

Strain KD145 (*ccna_03732*::*ccna_03732^A23T/G339N^* (*manA*^*^)) was cloned by PCR amplifying the *ccna_03732* locus from SDM1 genomic DNA with primers EK758/761. This fragment was ligated into the HindIII/EcoRI site of pNPTS138. The assembled plasmid (pEK227) was electroporated into NA1000 followed by selection on PYE-kanamycin plates. An individual colony was grown overnight in PYE and streaked onto PYE-3% sucrose plates. Colonies were screened by PCR amplifying the *ccna_03732* locus and DNA sequencing.

Strain KD175 (*ccna_03732*::*ccna_03732^A23T^*) was cloned by PCR amplifying the *ccna_03732* locus from NA1000 genomic DNA with primers EK758/761. This fragment was ligated into the HindIII/EcoRI site of pNPTS138. The A23T SNP was introduced by inverse-PCR using primers EK742/743. The assembled plasmid (pKD184) was electroporated into NA1000 followed by selection on PYE-kanamycin plates. An individual colony was grown overnight in PYE and streaked onto PYE-3% sucrose plates. Colonies were screened by amplifying genomic DNA with primers EKS188/740 and digesting with PpuMI (wild-type: 509 bp, A23T: 348 and 161 bp).

Strain KD174 (*ccna_03732*::*ccna_03732^G339N^*) was cloned by PCR amplifying the *ccna_03732* locus from NA1000 genomic DNA with primers EK758/761. This fragment was ligated into the HindIII/EcoRI site of pNPTS138. The G339N SNP was introduced by inverse-PCR using primers EK887/888. The assembled plasmid (pKD185) was electroporated into NA1000 followed by selection on PYE-kanamycin plates. An individual colony was grown overnight in PYE and streaked onto PYE-3% sucrose plates. Colonies were screened by amplifying genomic DNA with primers EKS191/Q42R and digesting with HpyCH4III (wild-type: 436 and 34 bp, G339N: 227, 209, and 34 bp).

Strains KD160 (*manA::manA-mCherry*), KD161 (*manA*^*^*::manA*^*^*-mCherry*), and KD162 (-13 ΔC *manA*^*^*::manA*^*^*-mCherry*) were cloned by Gibson assembly. The 5’-arm of *manA* was PCR amplified from NA1000 or *manA*^*^ genomic DNA with primers EK985/986; mCherry was amplified from pXCHYC-5 using primers EK987/988; the 3’-arm of *manA* was amplified from genomic DNA using primers EK989/990. Plasmid pNPTS138 was amplified with primers EK897/898. The assembled plasmids (pEK158 and pEK159) were electroporated into NA1000, *manA*^*^, or KD170 followed by selection on PYE-kanamycin plates. An individual colony was grown overnight in PYE and streaked onto PYE-3% sucrose plates. Colonies were screened by amplifying genomic DNA with primers EKS191/S65.

Strain KD186 (-13 ΔC *manA*^*^ promoter revertant) was cloned by Gibson assembly. The wild-type *manA* promoter was PCR amplified from *manA*^*^ genomic DNA with primers EK995/996. Plasmid pNPTS138 was amplified with primers EK897/898. The assembled plasmid (pKD165) was electroporated into KD170 followed by selection on PYE-kanamycin plates. An individual colony was grown overnight in PYE and streaked onto PYE-3% sucrose plates. Colonies were screened by amplifying genomic DNA with primers EKS191/759 and Sanger sequencing.

Strains KD64 (SDM1; P*xylX*-*ccna_03732*) and KD146 (*manA*^*^; P*xylX*-*ccna_03732*) were produced by electroporating plasmid pKD144 into SDM1 and *manA*^*^, respectively. Plasmid pKD144 was constructed by PCR-amplifying *ccna_03732* using primers EK739/740 and ligating into the NdeI/NheI site of pXYFPC-5.

Strain KD152 (SDM1; P*xylX*-*ccna_03732-FLAG*) was produced by electroporating plasmid pEK195 into SDM1. Plasmid pEK195 was constructed by PCR-amplifying *ccna_03732* using primers EK879/880 and ligating into the NdeI/NheI site of pXYFPC-5.

Strain KD153 (SDM1; P*xylX*-*yihS-FLAG*) was produced by electroporating plasmid pEK196 into SDM1. Plasmid pEK196 was constructed by PCR-amplifying *yihS* from *E. coli* genomic DNA using primers EK751/752 and ligating into the NdeI/NheI site of pXYFPC-5. A FLAG-tag was inserted by inverse-PCR of the resulting plasmid with primers EK817/818.

Strain EK266 (SDM1; P*xylX*-*rffE-FLAG*) was produced by electroporating plasmid pEK260 into SDM1. Plasmid pEK260 was constructed by PCR-amplifying *rffE* from *E. coli* genomic DNA using primers EK884/885 and ligating into the NdeI/NheI site of pXCHYC-5.

Strain EK270 (SDM1; P*xylX*-*nanE-FLAG*) was produced by electroporating plasmid pEK264 into SDM1. Plasmid pEK264 was constructed by Gibson assembly. *nanE* was PCR-amplified from *E. coli* genomic DNA using primers EK935/936 and pXCHYC-5 was amplified with primers EK921/922.

Strain EK268 (SDM1; P*xylX*-*manA-FLAG*) was produced by electroporating plasmid pEK263 into SDM1. Plasmid pEK263 was constructed by Gibson assembly. *manA* was PCR-amplified from *E. coli* genomic DNA using primers EK933/934 and pXCHYC-5 was amplified with primers EK921/922.

Strain KD181 (Δ*spdR*) was cloned by PCR amplifying the upstream (EK1081/1082) and downstream (EK1083/1084) homology fragments from NA1000 genomic DNA. The fragments were stitched together by overlap PCR. The final purified PCR product was ligated into the HindIII/EcoRI site of pNPTS138. The assembled plasmid (pKD176) was electroporated into NA1000 followed by selection on PYE-kanamycin plates. An individual colony was grown overnight in PYE and streaked onto PYE-3% sucrose plates. Colonies were screened for the *spdR* deletion with primers EK S247/S248 (wild-type 1.8 kb; deletion 1.4 kb).

Strain KD178 (*manA*^*^ + P*xylX*::*relA* (N-terminal domain)-*FLAG*) was produced by electroporating plasmid pKD187 into *manA*^*^. Plasmid pKD187 was constructed by PCR-amplifying the N-terminal domain of *relA* from *E. coli* MG1655 genomic DNA using primers EK1085/1086 and ligating into the NdeI/NheI site of pXCHYC-5.

**Table S1. Strains used in this study.**

| **Strain** | **Genotype** | **Construction** | **Source** |
| --- | --- | --- | --- |
| *C. crescentus* | | | |
| CB15 | Wild-type *C. crescentus* strain CB15 |  | (1) |
| NA1000 | Synchronizable variant of wild-type *C. crescentus* strain CB15 |  | (2) |
| KD80 | Stalk-deficient mutant-1 (SDM1) | NTG-mutagenesis screen | This study |
| KD9 | SDM1 + cosmid 2G12 | Conjugation of cosmid 2G12 into SDM1 | This study |
| KD10 | SDM1 + cosmid 2H1 | Conjugation of cosmid 2H1 into SDM1 | This study |
| KD64 | SDM1, P*xylX*::*ccna_03732* | Transformation of SDM1 with plasmid pKD144 | This study |
| KD145 | *ccna_03732*::*ccna_03732^A23T/G339N^* (*manA*^*^) (SDM2) | Transformation of NA1000 with plasmid pEK227 and sucrose selection | This study |
| KD146 | *manA*^*^, P*xylX*::*ccna_03732* | Transformation of *manA*^*^ with plasmid pKD144 | This study |
| KD152 | SDM1, P*xylX*::*ccna_03732-FLAG* | Transformation of SDM1 with plasmid pEK195 | This study |
| KD174 | *ccna_03732*::*ccna_03732^A23T^* | Transformation of NA1000 with plasmid pKD184 and sucrose selection | This study |
| KD175 | *ccna_03732*::*ccna_03732^G339N^* | Transformation of NA1000 with plasmid pKD185 and sucrose selection | This study |
| KD153 | SDM1 + P*xylX*::*yihS-FLAG* | Transformation of SDM1 with plasmid pEK196 | This study |
| EK203 | SDM1 + P*xylX*::*yihS-FLAG* (codon-optimized) | Transformation of SDM1 with plasmid pEK228 | This study |
| EK266 | SDM1 + P*xylX*::*rffE-FLAG* | Transformation of SDM1 with plasmid pEK260 | This study |
| EK270 | SDM1 + P*xylX*::*nanE-FLAG* | Transformation of SDM1 with plasmid pEK264 | This study |
| EK268 | SDM1 + P*xylX*::*manA-FLAG* | Transformation of SDM1 with plasmid pEK263 | This study |
| KD160 | *manA*::*manA-mCherry* | Transformation of NA1000 with plasmid pKD158 and sucrose selection | This study |
| KD161 | *manA*^*^::*manA*^*^*-mCherry* | Transformation of *manA*^*^ with plasmid pKD159 and sucrose selection | This study |
| KD170 | -13 ΔC *manA*^*^ promoter | Suppressor screen for recovery of stalk elongation | This study |
| KD162 | -13 ΔC *manA*^*^ promoter; *manA*^*^::*manA*^*^*-mCherry* | Transformation of strain KD170 with plasmid pKD159 and sucrose selection | This study |
| KD186 | -13 ΔC *manA*^*^ promoter revertant | Transformation of strain KD170 with plasmid pKD165 and sucrose selection | This study |
| EK717 | ΔMGE (mobile-genetic element) |  | (3) |
| CJW1249 | Tn5::*wbqP* |  | (4, 5) |
| KD181 | Δ*spdR* | Transformation of NA1000 with plasmid pEK176 and sucrose selection | This study |
| KD178 | *manA*^*^ + P*xylX*::*relA* (N-terminal domain)-*FLAG* | Transformation of *manA*^*^ with plasmid pKD187 | This study |
| *E. coli* | | | |
| S17-1 | λ−pir cloning strain |  | (6) |
| MG1655 | K-12 *E. coli* strain |  |  |
| 2G12 | *C. crescentus* genomic cosmid 2G12 |  | (7) |
| 2H1 | *C. crescentus* genomic cosmid 2H1 |  | (7) |

Table S2. Plasmids used in this study.

| **Name** | **Description** | **Source** |
| --- | --- | --- |
| pNPTS138 | *sacB*-containing suicide vector used for double homologous recombination, Kan^R^ | Alley, M.R.K. (unpublished) |
| pXCHYC-5 | Xylose-inducible expression, Tet^R^ | (8) |
| pKD144 | pXCHYC-5-based plasmid for *ccna_03732* expression, Tet^R^ | This study |
| pEK227 | pNPTS138-based plasmid to introduce *ccna_03732^A23T/G339N^* at the chromosomal locus, Kan^R^ | This study |
| pKD184 | pNPTS138-based plasmid to introduce *ccna_03732^A23T^* at the chromosomal locus, Kan^R^ | This study |
| pKD185 | pNPTS138-based plasmid to introduce *ccna_03732^G339N^* at the chromosomal locus, Kan^R^ | This study |
| pEK195 | pXCHYC-5-based plasmid for *ccna_03732-FLAG* expression, Tet^R^ | This study |
| pEK196 | pXCHYC-5-based plasmid for *yihS-FLAG* expression, Tet^R^ | This study |
| pEK228 | pXCHYC-5-based plasmid for *yihS-FLAG* expression, codon optimized, Tet^R^ | Synthesized by GenScript for this study |
| pEK260 | pXCHYC-5-based plasmid for *rffE-FLAG* expression, Tet^R^ | This study |
| pEK264 | pXCHYC-5-based plasmid for *nanE-FLAG* expression, Tet^R^ | This study |
| pEK263 | pXCHYC-5-based plasmid for *manA-FLAG* expression, Tet^R^ | This study |
| pKD158 | pNPTS138-based plasmid to introduce a C-terminal mCherry fusion at the chromosomal *manA* locus, Kan^R^ | This study |
| pKD159 | pNPTS138-based plasmid to introduce a C-terminal mCherry fusion at the chromosomal *manA*^*^ locus, Kan^R^ | This study |
| pKD165 | pNPTS138-based plasmid to introduce reintroduce the deleted -13 C in the *manA*^*^ suppressor strain, Kan^R^ | This study |
| pKD187 | pXCHYC-5-based plasmid for *relA* (N-terminal domain)-*FLAG* expression, Tet^R^ | This study |
| pKD176 | pNPTS138-based plasmid to delete *spdR*, Kan^R^ | This study |

**Table S3. Primers used in this study.**

| **Name** | **Sequence** |
| --- | --- |
| EK739 | tactcatATGACCAACGCTTTCGCCGA |
| EK740 | tacttctagaTCACGAAGCCGCGTTGATCA |
| EK740 | tacttctagaTCACGAAGCCGCGTTGATCA |
| EK742 | aCCTGATGGCCTATCTGCGCA |
| EK743 | CCTCAGCGGCGGCGGCGG |
| EK751 | tactcatatgAAATGGTTTAACACCCTAAGCCACAAC |
| EK752 | tactgctagcTTATTTCGCATTAATATCCAGCAGACCCGC |
| EK758 | tactaagcttAGGTCGCAGACCCCATTC |
| EK759 | GTTTCGTCGGACCGCCCAGGTCTTCAG |
| EK761 | tactgaattcACGGCTCGTCCAATGTTCT |
| EK817 | gactacaaggatgacgatgacaagTAAgctagctgcagcccgg |
| EK818 | TTTCGCATTAATATCCAGCAGACCCGC |
| EK879 | tactcatATGGAGCCTCCGATGACCAA |
| EK880 | tactgctagcTCActtgtcatcgtcatccttgtagtcCGAAGCCGCGTTGATCACC |
| EK884 | tactcatATGAAAGTACTGACTGTATTTGGTACGCG |
| EK885 | tactgctagcTCActtgtcatcgtcatccttgtagtcTAGTGATATCCGATTATTTTTTAACGCTTCCAGA |
| EK887 | GAAGACCTGGaCGGTCGAGGCG |
| EK888 | AGGCGGTCGCGAAGTCGG |
| EK897 | AAGCTTGGCGCCAGCCGG |
| EK898 | GAATTCGCTAGCTTCGGC |
| EK921 | GCTAGCTGCAGCCCGGGG |
| EK922 | ATGGTCGTCTCCCCAAAAC |
| EK933 | tggggagacgaccatATGCAAAAACTCATTAACTC |
| EK934 | cgggctgcagctagcTTActtgtcatcgtcatccttgtagtcCAGCTTGTTGTAAACAC |
| EK935 | tggggagacgaccatATGTCGTTACTTGCACAAC |
| EK936 | cgggctgcagctagcTTActtgtcatcgtcatccttgtagtcTAGCACCGCCTTTTTC |
| EK985 | agccggctggcgccaagcttTCCGTGAGTTCTTCGACC |
| EK986 | tcgaattctcCGAAGCCGCGTTGATCAC |
| EK987 | cgcggcttcgGAGAATTCGAACGTTACG |
| EK988 | accttgctggTTACTTGTACAGCTCGTC |
| EK989 | gtacaagtaaCCAGCAAGGTCGCCGCCC |
| EK990 | cggccgaagctagcgaattcCTGCGGCGTTCGCCTGGTC |
| EK995 | gctggcgccaagcttAGGTCGCAGACCCCATTC |
| EK996 | cgaagctagcgaattcTGCATGTGCGGATTGGAC |
| EK1081 | tactaagcttCAGGCCCGAGATCATTCA |
| EK1082 | accgcacatCAGCAGCAGAAGCGACTTATC |
| EK1083 | ctgctgctgATGTGCGGTCACAACGTCT |
| EK1084 | tactgaattcGCTCGATCATCAACCTGTCC |
| EK1085 | tactcatATGGTTGCGGTAAGAAGTGCAC |
| EK1086 | tactgctagcttatttatcatcatcatctttataatccatGCCCATCTGCAGCTGGTAGG |
| EKQ42R | CGATAGGCGGACAGGAAGTA |
| EKS65 | CTTGAAGCGCATGAACTCCTTGATG |
| EKS188 | GCCAGATCCGTGAGTTCTTC |
| EKS191 | GATCGTGAACATAGCCGTGA |
| EKS247 | AGCCTGGTTCAGCTGGTG |
| EKS248 | CTGTCGGTGCTGGTCAATAA |

| QPCR Primers | Forward | Reverse |
| --- | --- | --- |
| *cspD* | CAGGCGATATCTTCGTGCAT | CGACAACCAGTCCCTTGG |
| *katG* | CCGACCTCTATGTGCTGGTC | GACCCCAGTACAGCTCTTCG |
| *phoB* | ACCTCGACAAGGAGGGCTAC | ATCCAGTCGAGGATGACCAG |
| *pstC* | TGTCGGACGACATCATCAAC | GGCAGGACGACTTTCTTGAC |
| *rpoD* | CTCTATGCGATCAACAAGCG | ATAGGCCTTGAGGAACTCGC |
| *rpoH* | GAGAGCGAGTGGCAGGACT | CTCTTCCAGAAGCGACATCC |
| *spdR* | CGCATGCTGTTCTGGACAT | GTTGCCATAGCCCGTCAG |
